## Supplementary Figures for "Constitutive Activation and Oncogenicity Are Mediated by Loss of Helical Structure at the Cytosolic Boundary of the Thrombopoietin Receptor"

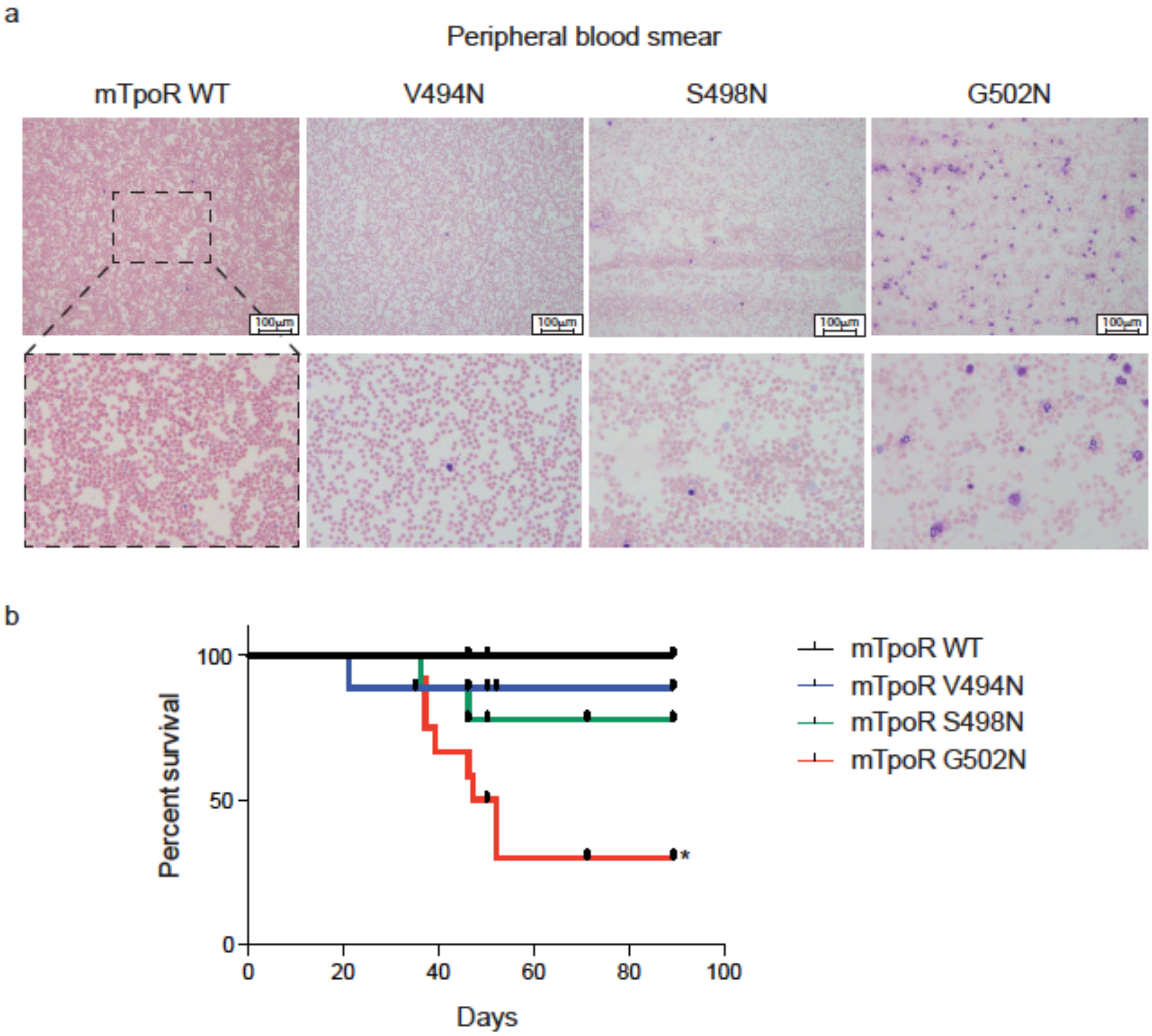


**Supplementary Figure 1. Mouse models expressing V494N, S498N and G502N mutants of TpoR show weak, mild and strong signs of myeloproliferation with a significant reduction of survival in the case of G502N mutants.** Peripheral blood smears from mouse models expressing the indicated constructs were stained with May-Grunwald Giemsa (10X). 3x-zoom was performed on the .jpg file using Adobe Illustrator (a). Kaplan-Meier survival analysis were performed on the indicated mice. The G502N model shows significant decrease of survival with a median survival of 49.5 days in the G502N mice. Statistical analysis was performed using the Mantel-Cox Test. *: $p<0.05$ (b).

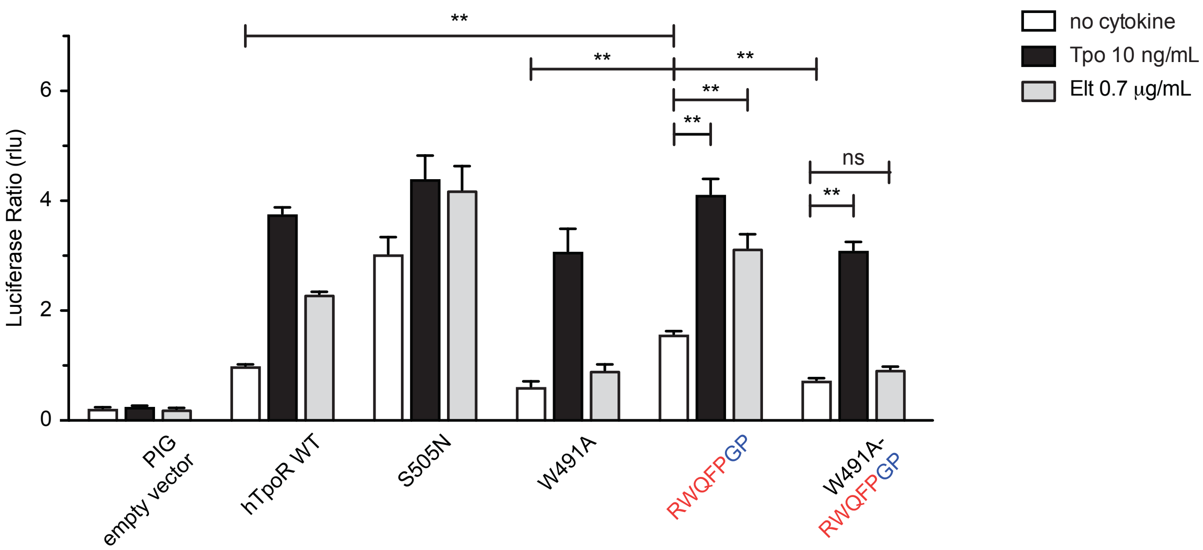


**Supplementary Figure 2. W491A mutation inhibits activation by Gly-Pro substitutions after the RWQFP motif.** STAT5 transcriptional activity reported by Spi-Luc in the absence or presence of thrombopoietin (Tpo) or eltrombopag (Elt) of human TpoR (hTpoR) wild type (WT) or mutants in HEK-239T cells. Shown are averages of three independent experiments each done with 3 biological replicates ± S.E.M. *: $p<0.05$, **: $p<0.01$, ns: non-significant; non-parametric multiple comparisons Steel's test with controls (jmp pro12).

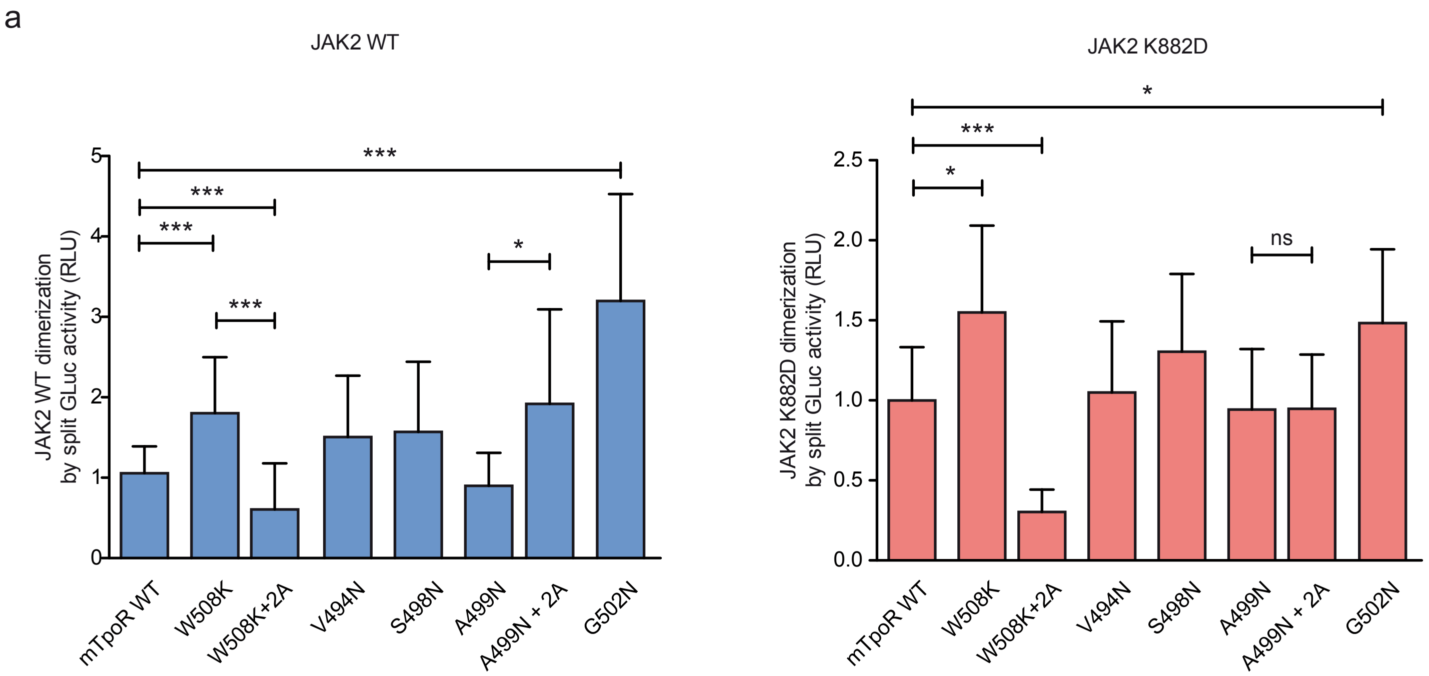


**Supplementary Figure 3. Dimerization of JAK2 is influenced by murine TpoR transmembrane mutations irrespectively of JAK2 activity.** Dimerization of JAK2 WT (a) and kinase-dead JAK2 K882D (b) bound to the indicated receptors was assessed by split Gaussia luciferase in HEK-293T cells (d, e, f). Shown are averages of separate experiments ± S.D. (n = 6 for JAK2 WT and n = 3 for JAK2 K882D); each experiment being performed with three biological repeats for each condition (triplicates). *: p<0.05, ***: p<0.001, ns: non-significant; non-parametric multiple comparisons Steel's test with controls.
